# Differential population declines across migratory strategies are shifting European bird communities towards residency

**DOI:** 10.64898/2026.11.18.767023

**Authors:** Antoine Havard, Etienne G. Henry, Inês Catry, Stanislas Rigal, Eva Šilarová, Jana Škorpilová, Dejan Bordjan, Stuart H. M. Butchart, Tomasz Chodkiewicz, Ingar Jostein Øien, Aleksi Lehikoinen, Åke Lindström, Bård G. Stokke, Sven Trautmann, Ana S.L. Rodrigues

**Affiliations:** CEFE, Univ Montpellier, CNRS, EPHE, IRD, Montpellier, France; CE3C Centre for Ecology, Evolution and Environmental Changes and CHANGE - Global Change and Sustainability Institute, Faculdade de Ciências da Universidade de Lisboa, 1749-016 Lisbon, Portugal; Foundation for Research on Biodiversity, Centre for the Synthesis and Analysis of Biodiversity, Montpellier, France; PECBMS, Czech Society for Ornithology, Prague, Czechia; Norwegian Institute for Nature Research, Trondheim, Norway; BirdLife International, David Attenborough Building, Pembroke Street, Cambridge CB2 3QZ, UK; Department of Zoology, Downing Street, Cambridge CB2 3EJ, UK; Museum & Institute of Zoology, Polish Academy of Sciences, Twarda 51/55, 00-818 Warszawa, Poland; Polish Society for the Protection of Birds (OTOP BirdLife Poland), Szeroki Dunaj 5, 00-255 Warszawa, Poland; BirdLife Norway, Sandgata 30b, NO-7012 Trondheim, Norway; Finnish Museum of Natural History, University of Helsinki, Finland; Department of Biology, Lund University, Sweden; DOPPS – DOPPS BirdLife Slovenia; Dachverband Deutscher Avifaunisten, An den Speichern 2, D-48149 Münster

**Keywords:** Migratory strategy, Population trends, Landbirds, Community changes

## Abstract

Previous studies revealed steep declines in populations of common migratory birds across Europe, especially for long-distance migrants. However, these studies focussed on continental-scale patterns, and they assigned a single migratory strategy to each species, often not distinguishing short-distance migrants from residents. It thus remains unclear whether the same patterns are found across European countries, and whether they hold when taking into account the variation in migratory behaviour observed across the European populations of many species. Here, we examine how European breeding landbird communities have varied according to migratory strategy between 1990 and 2023, at both continental and national scales. By compiling multiple sources of information on seasonal movements, we characterise the migratory strategy of landbird populations per country (long-distance migrant, short-distance migrant or resident). We combined these data with estimates of population trends and of abundance from national monitoring schemes to quantify changes in both multi-species indices (average change across species) and total abundance of birds, for each migratory strategy.

At the continental scale, multi-species indices reveal a strong decline in long-distance migrants (−26.8% between 1990 and 2023), whereas short-distance migrants declined more slowly (−4.5%) and residents remained stable. Changes in total abundance show even steeper declines – of 29.9% and 10.2% for, respectively, long- and short-distance migrants – whereas residents increased slightly (+3.4%). Together, these results point to a gradual reorganisation of bird communities towards a greater dominance of resident birds across Europe. However, these changes are not happening homogenously across the continent: whereas in Central and Southern Europe we observed the same general pattern of declining migrants while residents remained broadly stable or increased, in Northern Europe we reveal declines for all migratory strategies.

Overall, our study highlights the widespread vulnerability of migratory birds in Europe, while showing that the reorganisation of bird communities follows different trajectories across regions.

## Introduction

Bird communities worldwide are changing dramatically, including substantial recent declines documented across both Europe, where populations abundance have declined by 25% between 1980 and 2016 (Rigal et al. 2023), and North America, where they have declined by 29% between 1970 and 2018 (Rosenberg et al. 2019). The magnitude of these changes, however, is not uniform across taxa but structured according to species’ ecological traits. In Europe, for example, common species have declined more than rare ones (Inger et al. 2015), with species associated with farmland, cold environments, and urban areas having declined, respectively, by 57%, 40%, and 28% (Rigal et al., 2023). In contrast, habitat generalists increased by nearly 20% in France (Fontaine et al., 2021), while communities in both Europe and the United States are becoming more dominated by warm-dwelling species (Devictor et al., 2008; Stephens et al., 2016). Together, these patterns point to significant ongoing shifts in community composition and structure, driven primarily by anthropogenic changes including agricultural intensification (Doxa et al., 2010; Monnet et al., 2026; Rigal et al., 2023), habitat fragmentation (Lampila et al., 2005), and climate change (Lehikoinen et al. 2021).

Migration is a major ecological process structuring avian communities. Nearly one-fifth of all bird species undertake regular seasonal movements between spatially distinct breeding and non-breeding ranges (BirdLife International, 2026; Kirby et al., 2008), particularly prevalent among species that breed in the Northern Hemisphere. However, among species whose population trends have been quantified, there is ample evidence that migratory species in general (Marra et al., 2026), and long-distance migrants in particular, are declining disproportionately rapidly, a pattern observed in Europe (Burns et al. 2021; Vickery et al. 2023; Sanderson et al. 2006), North America (Rosenberg et al., 2019) and East Asia (Yong et al., 2021). This suggests there is an ongoing reorganisation of bird communities according to species’ migration strategy in response to global environmental change (Vickery et al., 2023).

In Europe, which shares billions of birds with Africa each year as they migrate along the African-Eurasian flyway (Newton, 2023), evidence of migratory bird declines comes from studies that analysed abundance and population trend data collected through national monitoring programmes. Despite using a variety of analytical approaches, and based on somewhat different sets of species and countries, these studies consistently found substantial declines in long-distance migrants over the past few decades (Burns et al. 2021; Vickery et al. 2023; Sanderson et al. 2006; Howard et al. 2020; Table 1). There is substantially more variation in the patterns of temporal change reported for shorter-distance migrants as well as for resident species, which were typically found to be declining less than long-distance migrants, or even stable or increasing (Table 1). *A priori*, short-distance migrants are expected to follow intermediate demographic trajectories between long-distance migrants and resident species, as observed in some regional-scale analyses (Kamp et al. 2021; Kuresoo et al. 2011, but see Tirozzi et al. 2021). Indeed, like long-distance migrants, they experience cumulative exposure to spatially-separated threats across their annual cycle (Buchan et al., 2023; La Sorte et al., 2018), but are spared from the risks faced when crossing the Sahara and while in sub-Saharan Africa. Similarly, short-distance migrants are also subject to phenological mismatches between their migration timing and changing environment conditions in the breeding grounds, but to a lesser extent than long-distance migrants (Cuchot et al., 2025; Møller et al., 2008; Robertson et al., 2024).

**Table 1.**
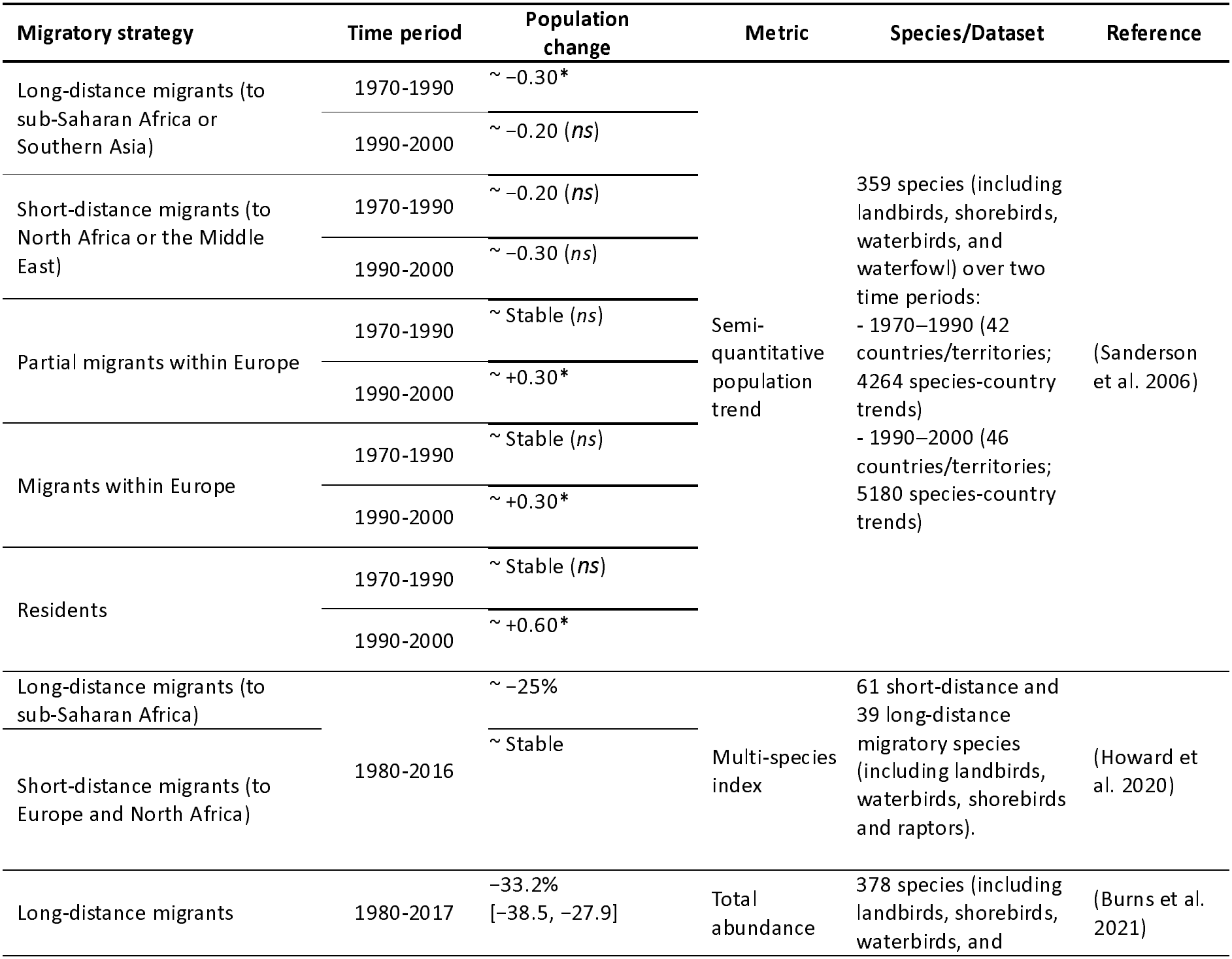
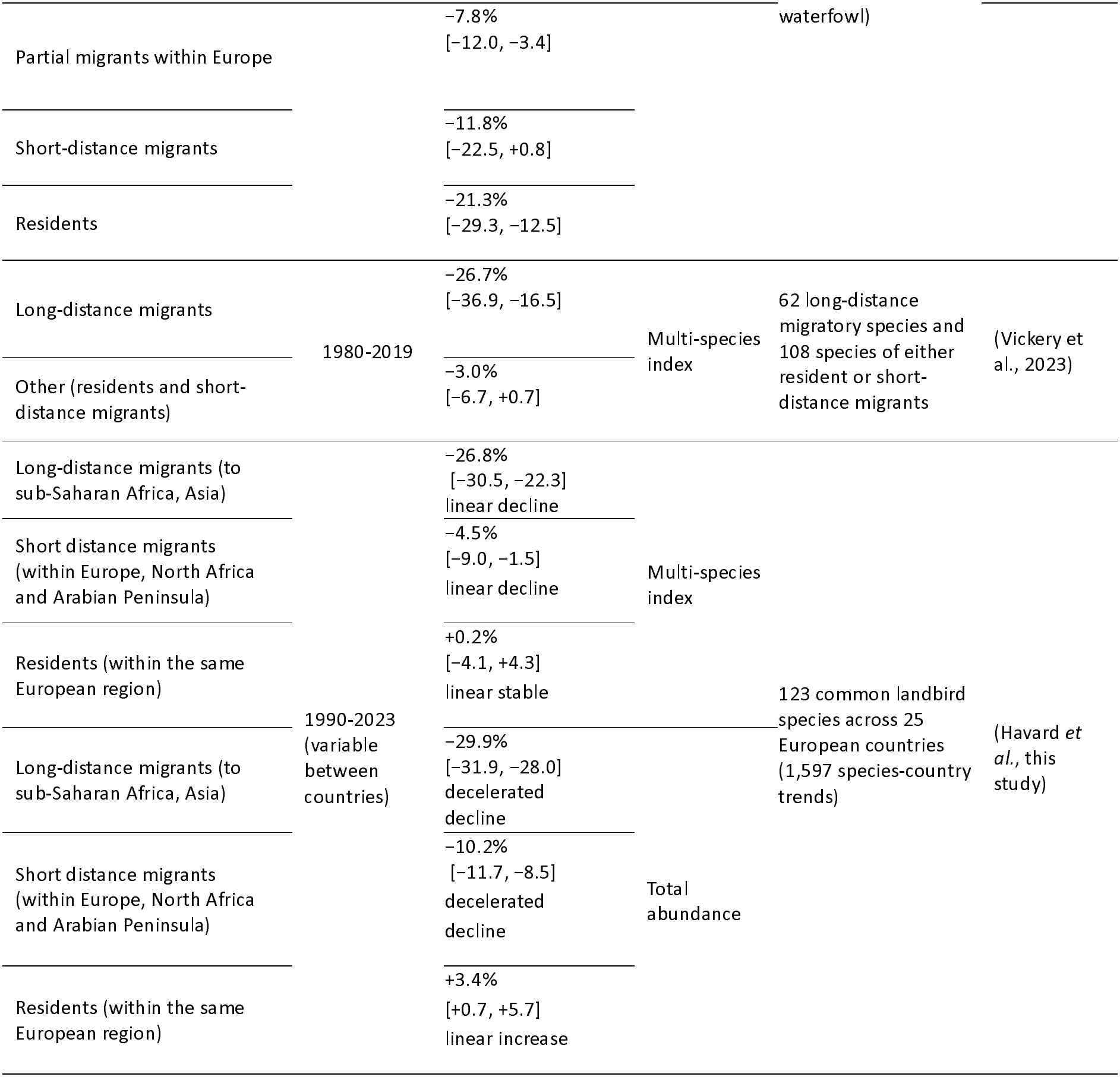
Overview of main studies that analysed multi-species trajectories in populations of birds according to migratory strategy at the European scale, including the present study. ‘ns’ indicates non-significant changes, * significant changes; ~ indicates approximate values estimated visually from published figures, where exact values were not reported in the text; values in square brackets indicate 95% confidence intervals (derived as estimate ± 1.96 × reported standard error when not available).

| Migratory strategy | Time period | Population change | Metric | Species/Dataset | Reference |
|---|---|---|---|---|---|
| Long-distance migrants (to sub-Saharan Africa or Southern Asia) | 1970-1990 | ~ −0.30* | Semi-quantitative population trend | 359 species (including landbirds, shorebirds, waterbirds, and waterfowl) over two time periods:<br>- 1970–1990 (42 countries/territories; 4264 species-country trends)<br>- 1990–2000 (46 countries/territories; 5180 species-country trends) | (Sanderson et al. 2006) |
|  | 1990-2000 | ~ −0.20 (*ns*) |  |  |  |
| Short-distance migrants (to North Africa or the Middle East) | 1970-1990 | ~ −0.20 (*ns*) |  |  |  |
|  | 1990-2000 | ~ −0.30 (*ns*) |  |  |  |
| Partial migrants within Europe | 1970-1990 | ~ Stable (*ns*) |  |  |  |
|  | 1990-2000 | ~ +0.30* |  |  |  |
| Migrants within Europe | 1970-1990 | ~ Stable (*ns*) |  |  |  |
|  | 1990-2000 | ~ +0.30* |  |  |  |
| Residents | 1970-1990 | ~ Stable (*ns*) |  |  |  |
|  | 1990-2000 | ~ +0.60* |  |  |  |
| Long-distance migrants (to sub-Saharan Africa) | 1980-2016 | ~ −25% | Multi-species index | 61 short-distance and 39 long-distance migratory species (including landbirds, waterbirds, shorebirds and raptors). | (Howard et al. 2020) |
| Short-distance migrants (to Europe and North Africa) |  | ~ Stable |  |  |  |
| Long-distance migrants | 1980-2017 | −33.2%<br>[−38.5, −27.9] | Total abundance | 378 species (including landbirds, shorebirds, waterbirds, and waterfowl) | (Burns et al. 2021) |
| Partial migrants within Europe |  | −7.8%<br>[−12.0, −3.4] |  |  |  |
| Short-distance migrants |  | −11.8%<br>[−22.5, +0.8] |  |  |  |
| Residents |  | −21.3%<br>[−29.3, −12.5] |  |  |  |
| Long-distance migrants | 1980-2019 | −26.7%<br>[−36.9, −16.5] | Multi-species index | 62 long-distance migratory species and 108 species of either resident or short-distance migrants | (Vickery et al., 2023) |
| Other (residents and short-distance migrants) |  | −3.0%<br>[−6.7, +0.7] |  |  |  |
| Long-distance migrants (to sub-Saharan Africa, Asia) | 1990-2023 (variable between countries) | −26.8%<br>[−30.5, −22.3]<br>linear decline | Multi-species index | 123 common landbird species across 25 European countries (1,597 species-country trends) | (Havard *et al.*, this study) |
| Short distance migrants (within Europe, North Africa and Arabian Peninsula) |  | −4.5%<br>[−9.0, −1.5]<br>linear decline |  |  |  |
| Residents (within the same European region) |  | +0.2%<br>[−4.1, +4.3]<br>linear stable |  |  |  |
| Long-distance migrants (to sub-Saharan Africa, Asia) |  | −29.9%<br>[−31.9, −28.0]<br>decelerated decline | Total abundance |  |  |
| Short distance migrants (within Europe, North Africa and Arabian Peninsula) |  | −10.2%<br>[−11.7, −8.5]<br>decelerated decline |  |  |  |
| Residents (within the same European region) |  | +3.4%<br>[+0.7, +5.7]<br>linear increase |  |  |  |

Previous studies investigating the relationship between migratory strategy and demographic trends at the European scale assigned a single category of migratory strategy (resident, short or long-distance migrant) to each species (Table 1). This is often an oversimplification, because there can be substantial variation in the migratory propensity of individuals across populations of the same species. For example, most Eurasian Wren *Troglodytes troglodytes* breeding in Sweden spend winter in Central Europe, whereas their English counterparts are largely resident (Kroodsma et al., 2020). Variation can even occur within populations: partially migratory populations comprise both resident and migratory individuals (Chapman et al., 2011). In Denmark, for example, most Eurasian Blackbirds (*Turdus merula)* are resident, but some winter in Britain (Collar et Christie 2020). Because individuals with different migratory strategies are exposed to different pressures throughout their life-cycle, analyses based on the assignation of a single migratory category to each species may obscure important demographic heterogeneity, potentially biasing continental-scale population trends. A finer population-level characterisation of migratory strategy is thus warranted to support more precise analyses of the relationship between migratory strategy and community changes.

Another gap in understanding how European bird communities are changing in relation to migratory strategies relates to the spatial variation across the continent. Indeed, the broad pattern observed at the European scale, of declining long-distance migrants and more stable resident and short-distance species (Howard et al. 2020; Vickery et al. 2023; Sanderson et al. 2006), is not necessarily found across all regions, both because of the above-mentioned inter- and intra-population variability in migratory propensity, and because of spatial distribution and prevalence of migratory species in bird communities. For example, bird communities in Northern Europe are more strongly dominated by migratory species due to stronger seasonality (Somveille et al. 2013), and may thus be experiencing greater levels of restructuring than those in southern regions. Moreover, the anthropogenic activities driving bird demographic changes also vary across space. In particular, recent agricultural intensification and urbanisation are particularly pronounced in Western Europe (Kuemmerle et al., 2016; Rigal et al., 2023), whereas recent temperature increases have been stronger toward Eastern and Northern Europe (Copernicus Climate Change Service, 2023). The fact that migratory species are exposed to environmental change across multiple regions further confounds the spatial patterns of demographic change observed across breeding populations. For example, western populations of the Eurasian Blackcap (*Sylvia atricapilla*) predominantly winter in Western Europe and the Western Mediterranean, whereas more eastern populations migrate toward South-Eastern Europe and Eastern Africa (Delmore et al., 2020). This pattern reflects a broader organisation of migratory connectivity observed across European migratory bird species at flyway scale (Briedis et al., 2020), resulting in population-specific exposure to environmental change (Carboneras et al., 2024). Such variation in migratory routes and wintering destinations may result in different population dynamics (Runge et al., 2014), and may help to explain the spatial heterogeneity in demographic trends observed across Europe among breeding species with different migratory strategies (Bowler et al. 2021).

Finally, previous studies have quantified population changes using one of two types of metrics. Some used multi-species indices, calculated by averaging changes in relative population sizes across species (Howard et al. 2020; Vickery et al. 2023). Others accounted for differences in absolute population size, so that their results reflect changes in the total numbers of individuals (Sanderson et al. 2006; Burns et al. 2021). Both types of metrics are informative, as change in a multi-species index reflects the average response of the community, to which rare and common species contribute equally, whereas change in total abundance is primarily driven by the most numerous species, which have often a higher weight in driving ecosystem functioning (Gaston 2010; Inger et al. 2015). Consequently, these two metrics may lead to different conclusions if abundant species and rarer species respond differently; this, combined with variations in the set of species, countries and time periods covered, limits the comparability across different studies (Table 1).

In this study, we present a more complete picture of the ongoing changes in European breeding landbird communities in relation to migratory strategies by addressing these limitations of previous research: we take into account intraspecific variation in migratory strategies, through an approach that spans regional to continental scales, contrasting different metrics of change. First, we characterise the national populations of each of 123 species of European common birds according to their migratory strategies, by taking into account inter-as well as intra-population variability in migratory propensity. Then, we integrate this information with national data on the abundance and population trends of these populations to quantify separately the temporal dynamics of bird populations of residents, short-distance migrants, and long-distance migrants, at both the national level and across the European continent. By deriving both multi-species indices and trends in total abundances for each of these migratory strategies, we provide a standardised and comprehensive assessment of population dynamics in relation to migratory strategy across Europe.

We predict that communities are reorganising towards an increasing prevalence of residents relative to migrants, which should be observable both through multi-species indices and in temporal series of total abundance. We further predict that long-distance migrants have declined the most, and residents the least, with short-distance migrants showing intermediate trends. Finally, given that communities in northern countries are more strongly dominated by migratory birds, we predict them to be more strongly affected by these changes.

## Methods

Full methodological details are provided in Supplementary Methods, here we present an overview.

This study focuses on common European breeding landbirds, comprising non-raptor species that rely primarily on terrestrial habitats for both feeding and breeding, thereby excluding waterbirds and seabirds. The ecological units considered are “national populations”, defined as all individuals of a given species breeding within a given country (e.g., Dunnocks *Prunella modularis* breeding in Switzerland). We included in this analysis species (Table S1) and national populations (Table S2) for which data on population trends, population size, and migratory strategy were simultaneously available.

### Migratory strategies of national populations or sub-populations

We assigned migratory strategies to national populations based on information on their distribution outside the breeding season, excluding migratory periods (herein, ‘wintering’, for simplicity) across the 15 biogeographic regions used in the Eurasian–African Bird Migration Atlas (EABMA; Figure S1). We considered three strategies: (i) ‘resident’, wintering in the same European region; (ii) ‘short-distance migrant’, wintering in another European region, North Africa or the Arabian Peninsula; and (iii) ‘long-distance migrant’, migrating to sub-Saharan Africa or tropical Asia.

We derived information on wintering distributions primarily from Fattorini et al. (2023), who used ringing and recovery data compiled by the European Union for Bird Ringing (EURING; Du Feu et al. 2016) as part of the EABMA (Spina et al. 2022). For 125 European breeding species, they measured the proportion of individuals ringed in a given breeding region that were subsequently re-encountered in each region during the non-breeding season. We corrected these proportions by considering variation in ringing effort (Figure S2) to obtain estimates of the fraction of individuals in each breeding population that follow each of the three migratory strategies. National populations containing both resident and migratory individuals were considered partially migratory and subdivided into sub-populations.

Fattorini et al. (2023) mainly covered resident and short-distance migratory populations. We used seasonal distribution maps from BirdLife International and Handbook of the Birds of the World (2024) to classify 49 additional species as long-distance migrants (with non-breeding range fully outside Europe and North Africa), and three others as short-distance migrants (with non-overlapping breeding and non-breeding distributions entirely north of the Sahara). Finally, we classified as resident 21 species based on descriptions of predominantly sedentary behaviour across Europe in Birds of the World (Billerman et al., 2025). We excluded from the analysis nine species for which we could not reliably assign a migratory strategy per population (e.g., the Woodlark *Lullula arborea*, which has partially migratory populations but is not covered by the EABMA; Table S1).

### Population and sub-population abundance time series

We obtained national annual population index time series from the Pan-European Common Bird Monitoring Scheme (PECBMS; https://pecbms.info/; 30 European countries), which reflect relative changes in national population size. We retained data for 25 European countries whose monitoring schemes covered at least 15 years between 1990 and 2023 (Table S3). In addition, we obtained national population size estimates from the European Red List of Birds (ERLoB; BirdLife International 2021). We then combined these two sources to derive abundance time series (i.e. number of individuals per year) for each national population, following Rigal et al. (2023). In the case of populations with mixed migratory strategies, we divided the total population size among sub-populations according to the proportion of individuals assigned to each migratory strategy, and we assumed similar population dynamics for each sub-population.

We obtained a final dataset comprising 2,087 national populations and sub-populations, corresponding to 123 species across 25 European countries. This dataset had excellent coverage (>90%) of total landbird abundance for 18 out of the 25 countries analysed, and a good coverage (75– 90%) for four others, with three countries having lower coverage (Greece, 60.5%; Bulgaria, 54.1%; Romania, 41.8%; Figure S3).

### National trends per migratory strategy: multi-species indices and total abundance

We excluded Greece, Romania and Bulgaria from the national-scale analyses, given the poorer coverage of our dataset for these countries (Figure S3). For each of the 22 countries analysed, we aggregated all populations and sub-populations sharing the same migratory strategy to derive two complementary metrics of temporal change (i.e., time series) per migratory strategy. The first is a multi-species index, expressed annually in relation to a 1990 baseline, corresponding to a mean change in abundance where species are given equal weight. The second metric estimates the total abundance (i.e., number of individual birds) per year, with contributing species weighted based on their absolute population size (Rigal et al. 2023). For both metrics, we followed Soldaat et al. (2017) to derive geometric means and 90% confidence intervals (through Monte Carlo simulations) for each national time series, for each migratory strategy.

To allow comparisons between countries with differing monitoring durations, we standardised percentage changes by dividing them by the number of monitored years, yielding annualised (henceforth “annual”) percentages of change. We mapped these to visualise the spatial patterns in multi-species indices and total abundance dynamics across Europe

### European trends per migratory strategy: multi-species indices and total abundance

We obtained European annual trends using the same two metrics – multi-species indices and total abundance – for each migratory strategy by aggregating national species time series, following Pannekoek et van Strien (2005) and Rigal et al. (2023), using the rtrim R package (Bogaart et al., 2026), with each country weighted by population size. We excluded Spain from this continental-scale analysis because its national population sizes appear to be overestimated (Supplementary Methods; Figures S4 and S5).

### Characterising trends

We characterised each of the temporal trends obtained (i.e., multi-species indices and trends in total abundance, at the national and at the European scales, for each migratory strategy) in terms of the trajectory shape and the significance of the change. We followed Rigal et al. (2020) by modelling trajectories using second-degree polynomial regressions (allowing both linear and non-linear dynamics to be captured) while estimating confidence intervals. We used the confidence intervals to characterize each trend according to its overall direction (increase, decline, or stable) and the significance of the second-degree polynomial coefficients of the fitted tend to describe its shape (linear, accelerated, decelerated, convex, or concave; Table 2).

**Table 2.** Trend characterization according to the limits of the confidence interval and the significance of the second-degree polynomial coefficient (α_2_) of the fitted trend.

|  |  | Shape of trajectory |  |  |
|---|---|---|---|---|
|  | Second polynomial coefficient | α<sub>2</sub> *ns* | α<sub>2</sub> > 0 * | α<sub>2</sub> < 0 * |
| Confidence intervals | Above zero: increase | Linear increase | Accelerated increase | Decelerated increase |
|  | Crosses zero: stable | Linear stable | Convex stable | Concave stable |
|  | Below zero: decline | Linear decline | Decelerated decline | Accelerated decline |

### Contribution of each migratory strategy to total abundance

For each of the 22 analysed countries, we estimated the relative contribution of each migratory strategy (i.e. the percentage of individuals adopting the strategy, in relation to the total abundance of all analysed landbirds), for year 2023, from the above-mentioned estimates of total abundance per year.

### Changes in the prevalence of migratory birds

We estimated for each country the percentage change in the proportion of migrants (i.e., long- and short-distance migrants) relative to the abundance of all birds, again using a bootstrap procedure to propagate uncertainty. As for the national trends per migratory strategy, we express this as a percentage of change per year, characterising the corresponding changes in migratory prevalence.

### Changes in community composition per country

We also estimated the change in each population and subpopulation per country, by modelling abundance over time (using a similar bootstrap approach) and then contrasting abundance in the last versus the first year of monitoring.

### Sensitivity analyses

We examined the sensitivity of the results to three methodological choices. First, in our analysis we considered that France belongs in its entirety to West Europe (even though in the EABMA it was split into three regions; Figure S1), so we evaluated the effect of allocating instead France to South-west Europe. Second, we controlled for variation in ringing effort when calculating the transition rates between breeding and wintering regions, so we present an alternative analysis without this correction. We investigated the effects of these two choices both on the European-scale trends and in the national-scale patterns for short-distance migrants and resident species (these two choices have almost no effect on long-distance migrants). Third, in our analyses of trends at the European-scale we excluded Spain given its strong effect on the results, so as a sensitivity analysis we also present the results including Spain.

All analyses were performed in R version 4.3.2 (R Core Team, 2023).

## Results

### Contribution of different migratory strategies to national bird communities

We assessed the migratory strategy and demographic trends of 2,087 national populations (i.e., set of individuals of a given species breeding in a given country) or sub-populations (subset of a national population comprising individuals with the same migratory strategy), representing 123 species across 25 countries. Among these, 28% were classified as long-distance migrants, 29% as short-distance migrants and 43% as residents.

We found wide variation in the relative contribution of different migratory strategies to national breeding bird communities across Europe (Figure 1). Northern communities are dominated by migrants, with short- and long-distance migrants accounting for more than 75% of the landbird abundance in Fennoscandia, Latvia and Estonia. These regions also stand out for their prevalence of long-distance migrants (e.g., 43.5% in Estonia, 36.6% in Sweden).

**Figure 1.**
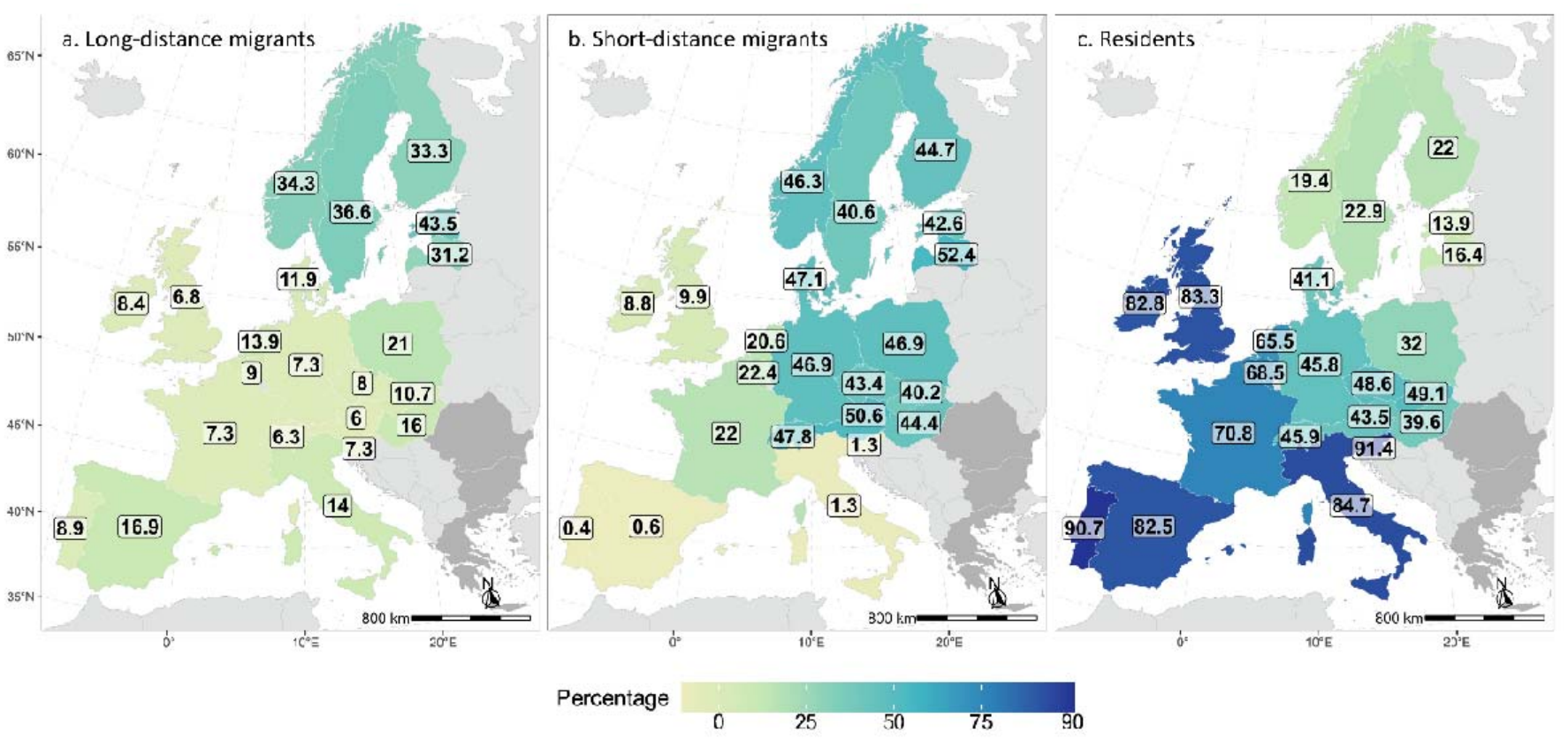
Relative contribution (in % of individual birds) of each migratory strategy to the total landbird abundance in each country for the year 2023: (a) long-distance migrants, (b) short-distance migrants, and (c) residents. Countries not included in the analysis are shown in grey.

The proportion of migrants decreases progressively towards lower latitudes. In Central Europe and Denmark, communities are dominated by residents and short-distance migrants in approximately equal proportions (e.g., 48.6% residents and 43.4% short-distance migrants in the Czech Republic). In Western Europe, communities are dominated by residents, although short-distance migrants still account for at least 20% of the total abundance (e.g., 22.0% in France).

Residents dominate even more strongly in the British Isles, representing about 83% of individuals, and in Southern European countries, where they account for more than 80% of individuals (e.g., 90.7% in Portugal and 84.7% in Italy). These southern countries correspondingly have the lowest proportions of short-distance migrants (generally below 3%), while long-distance migrants represent a larger share of communities, reaching 16.9% in Spain.

### European-scale trends

Across all analysed species, the European landbird community declined by 14.5% between 1990 and 2023 as measured through the multi-species index ([−11.2, −17.2]; linear decline) and by 9.2% in total abundance ([−10.4, −7.9]; decelerated decline) (Figure S6).

Long-distance migrants showed the strongest declines, with a 26.8% decrease in the multi-species index (95% CI = [−30.5, −22.3]; linear decline; Figure 2a) and a 29.9% decline in total abundance ([−31.9, −28.0]; decelerated decline) (Figure 2b). Short-distance migrants declined more moderately, losing 4.5% in terms of their multi-species index ([−9.0, −1.5]; linear decline) and 10.2% in terms of total abundance ([−11.7, −8.7]; decelerated decline). Residents, in contrast, remained stable when measured through the multi-species index (+0.2% [−4.1, +4.3]; linear stable) and increased slightly in total abundance (+3.4% [+0.7, +6.1]; linear increase).

**Figure 2.**
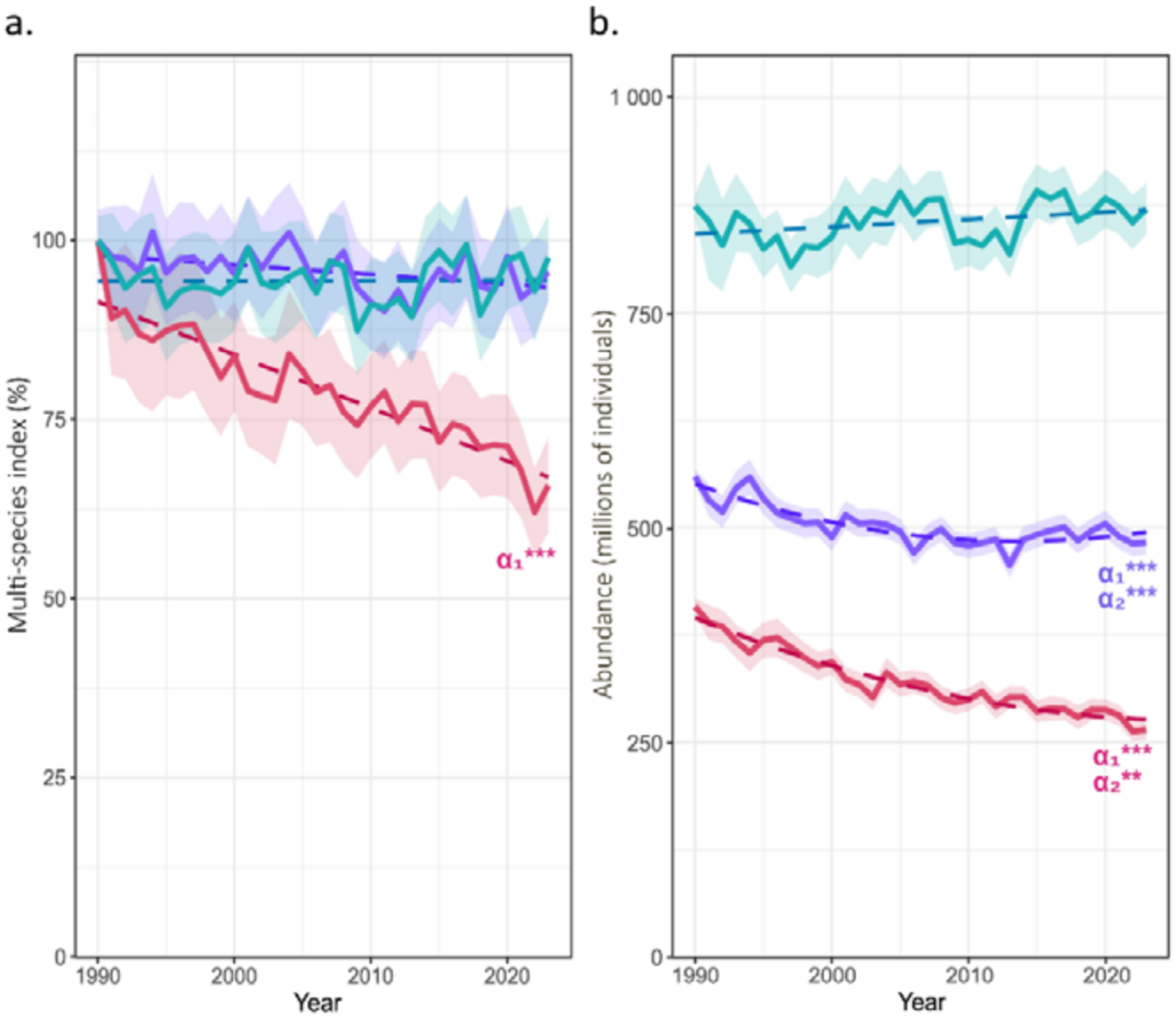
Temporal changes in landbird populations by migratory strategy across 24 European countries (excluding Spain), 1990–2023: (a) multi-species index (in relation to 1990, set to 100) and (b) total abundance (number of individuals). Solid lines represent the values for long-distance migrants (n = 51 species, red), short-distance migrants (n = 46 species, purple), and residents (n = 70 species, green). Shaded bands represent the corresponding 95% confidence intervals around the index values, and dotted lines show fitted polynomial trends. Asterisks (*, **, ***) show the significance levels (p < 0.05, p < 0.01, and p < 0.001, respectively) associated with the polynomial coefficients (α_1_ and α_2_) of the fitted trend.

Overall, the total abundance of migratory breeding individuals declined by 18.4% ([−17.0, −19.9]) over the study period.

### National-scale trends

National trajectories of the multi-species index varied substantially across countries and across migratory strategies (Figures 3, S7, Table S4). Of the 22 countries analysed, 16 experienced declines in their overall landbird communities, although the shape of these trends differed (Figure 3a). Nine countries showed linear declines, with Slovakia experiencing the steepest annual decline (−1.1% yr^−1^), while four exhibited decelerated declines (such as Germany and Belgium) and three accelerated declines (such as France or Czech Republic). Only four countries showed increases, with linear changes of +0.8% yr^−1^ in Hungary and +0.6% yr^−1^ in Switzerland.

**Figure 3.**
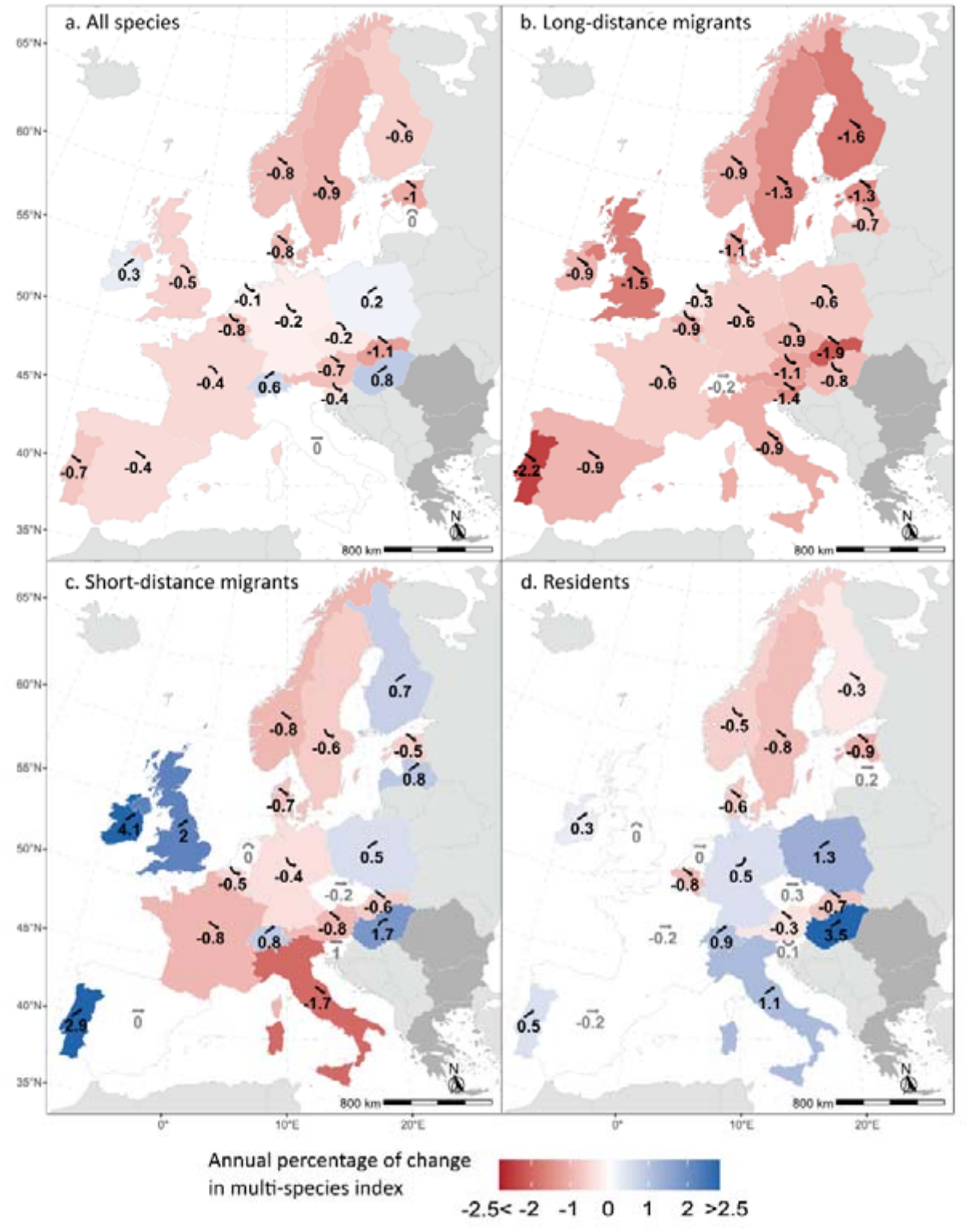
Annual percentage of change in multi-species index for (a) all landbirds, combining all migratory strategies, (b) long-distance migrants, (c) short-distance migrants, and (d) residents, in each of 22 European countries. Colours indicate the direction and magnitude of change: blue for significant increases, red for significant declines, white for no significant change. Symbols above each value represent the trajectory type (Table 2). Countries shown in dark grey were excluded from the country-level analyses.

The multi-species index of long-distance migrants declined across all European regions, in 21 of the 22 countries (Figure 3b): 13 showed linear declines, four decelerated and four accelerated. The steepest decline was observed in Portugal (−2.2% yr^−1^). Switzerland was the only exception, with a linear stable trajectory. Notably, no country showed an increase.

Short-distance migrants showed the most variable responses among countries (Figure 3c). Their index declined in ten countries (seven linear, three decelerated), increased in eight (seven linear, one decelerated), and was stable in the remaining four. The strongest increases occurred in Atlantic countries: Ireland (+4.1% yr^−1^), Portugal (+2.9% yr^−1^) and United Kingdom (+2.0% yr^−1^). In contrast, the steepest decline occurred in Italy (−1.7% yr^−1^).

The multi-species index of residents increased or remained stable in most countries (Figure 3d): it increased in seven countries (six linear, one accelerated), and was stable in seven (five linear, one concave, one convex). The most positive trends occurred in Central, South-Central, and South-Eastern Europe, reaching up to +3.5% yr^−1^ in Hungary. Declines were observed in eight countries (all linear except a country with a decelerated decline), mainly in northern countries (e.g., Estonia −0.9% yr^−1^).

Patterns of change in total abundance across countries (Figures 4, S8, Table S5) generally mirror those in the multi-species index, with significant positive Pearson correlations coefficient between the two metrics for all species (R^2^ = 0.56), long-distance migrants (R^2^ = 0.27), short-distance migrants (R^2^ = 0.33) and residents (R^2^ = 0.40).

**Figure 4.**
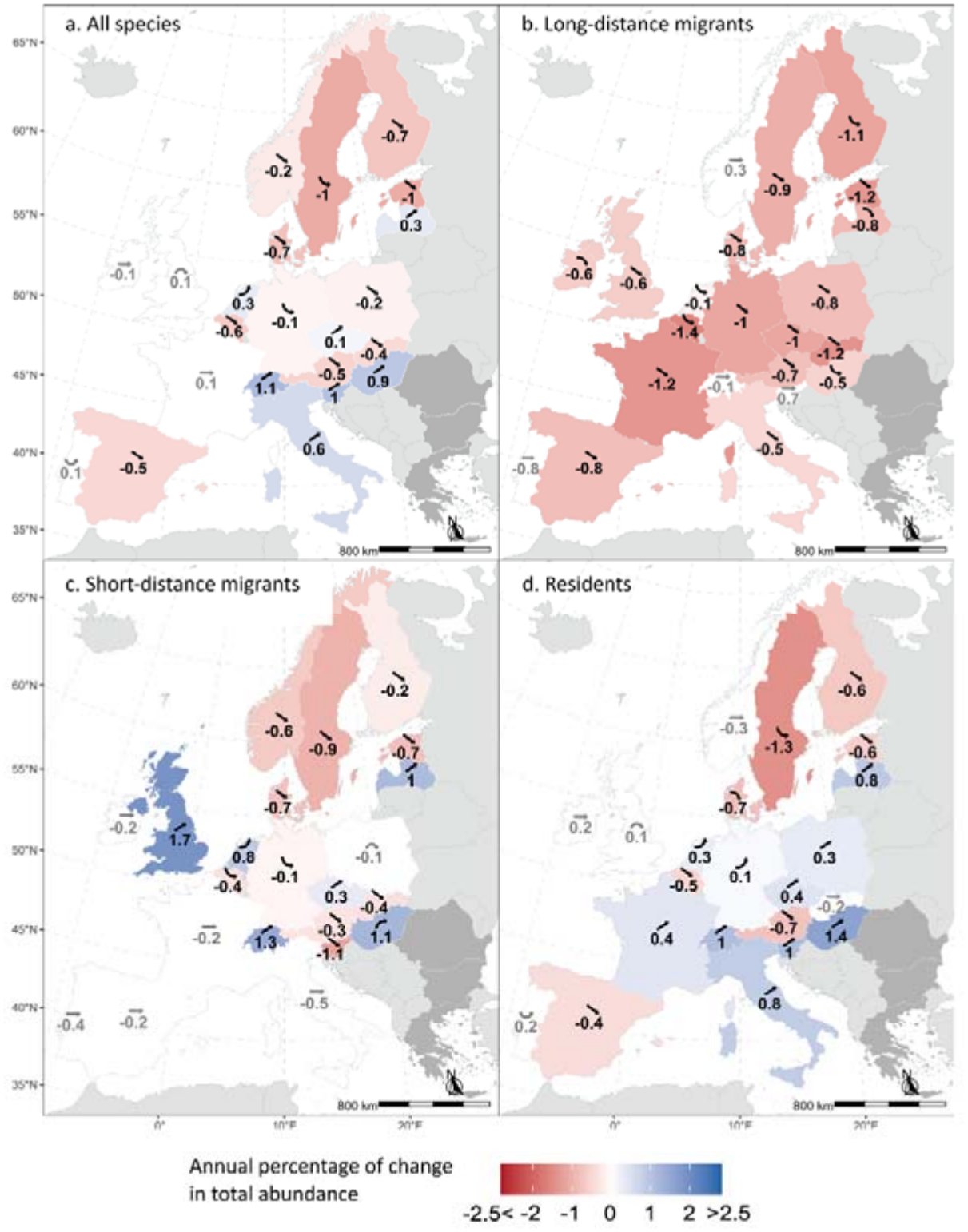
Annual percentage of change in total abundance for (a) all landbirds, combining all migratory strategies, (b) long-distance migrants, (c) short-distance migrants, and (d) residents, in each of 22 European countries. Colours indicate the direction and magnitude of change: blue for significant increases, red for significant declines, white for no significant change. Symbols above each value represent the trajectory type (Table 2). Countries shown in dark grey were excluded from the country-level analyses.

Across all species, total abundance declined in 11 countries (nine linear, two decelerated), with the strongest declines in Estonia and Sweden (−1.0% yr^−1^, Figure 4a). We found increases in seven countries (six linear and one accelerated), most strongly in Switzerland (+1.1% yr^−1^) and stable trajectories in four (two linear, one convex, one concave).

As in the multi-species index, nearly all countries (18 out of 22) showed declines in long-distance migrants (12 linear, four decelerated, two accelerated; Figure 4b), the steepest being observed in Belgium (−1.4% yr^−1^). No country showed increasing trends, while the remaining four countries exhibited stable linear trajectories.

Short-distance migrants again showed contrasting patterns among countries (Figure 4c), with a relative balance between declining (ten; eight linear, two decelerated), increasing (six; four linear, one accelerated, one decelerated) and stable trajectories (six; five linear, one concave). The steepest declines were again observed in Northern Europe (reaching −0.9% yr^−1^ in Sweden) whereas the strongest increase was found in the United Kingdom (+1.7% yr^−1^).

Total abundance of residents increased in ten countries (eight linear, two accelerated), up to +1.4% yr^−1^ in Hungary, and remained stable in five (three linear, one concave, one convex; Figure 4d). We found declining trajectories in seven countries (five linear, one accelerated, one decelerated), mostly in Northern Europe, the steepest being in Sweden (−1.3% yr^−1^; Figure 4d).

The trajectories of these national-scale indices are driven by variations (positive and negative) in the abundances of common species, that vary from country to country (Figure S9).

### Changes in the prevalence of migratory birds

The overall proportion of individual migratory birds (long- and short-distance migrants combined) within national breeding communities declined in 11 of the 22 countries analysed (Figure 5). The strongest decreases occurred in South-Central Europe, reaching −0.8% yr^−1^ in Italy, and gradually weakening toward Central Europe (e.g. −0.2% yr^−1^ in Germany). Austria was the only Central European country where the proportion of migrants increased (linear increase, +0.2% yr^−1^).

**Figure 5.**
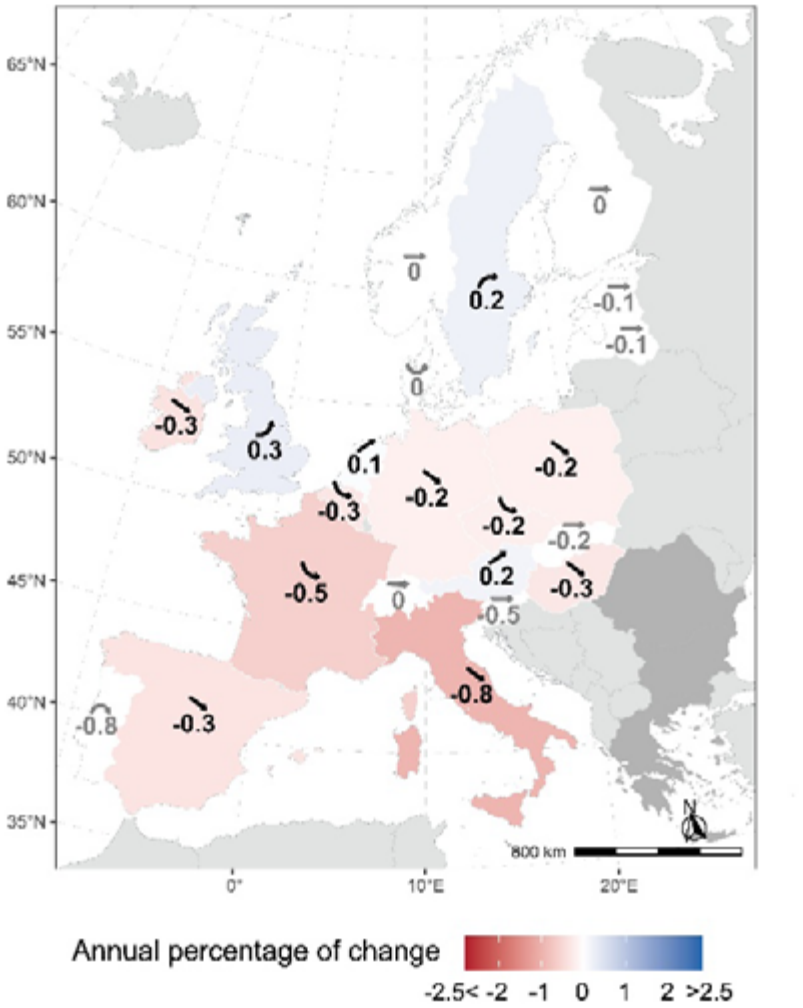
Annual change in the number of individual migratory birds as a proportion of total breeding landbird abundance across European countries. Colours indicate the direction and magnitude of change. Symbols above each value represent the trajectory type (table 2). Countries shown in dark grey were excluded from the country-level analyses.

In contrast, northern countries generally exhibit stable or even increasing trends in the proportion of migratory birds, with an accelerated increase in the United Kingdom (+0.3% yr^−1^), a decelerated increase in Sweden (+0.2% yr^−1^) and stable trajectories in other Fennoscandian countries. Across Europe, the proportion of migrants declined by −11.4% (decelerated decline).

### Sensitivity analyses

Results are robust to the methodological choices tested through sensitivity analyses.

At the European scale, we obtained qualitatively similar results when including Spain in the analysis, the main difference being that, in that case the trends for resident species show a significant increase in absolute numbers (Figure S10).

The results were also robust to the methodological choices of considering France wholly within West Europe (and not within South-west Europe; Figures S11, S12, S13) and to correcting species transitions rates across regions based on ringing effort (rather than not correcting at all; Figure S14, S15, S16). Indeed, broadly similar results were obtained for European-scale trends (Figures S11, S14) and in terms of spatial patterns across European countries (Figures S12, S13, S15, S16), even if some national estimates are modified for short-distance migrants and residents, particularly in southern countries.

## Discussion

### A new dataset linking population-level bird migratory behaviour to demographic changes

We characterised migratory behaviour at the population level and linked it to changes in bird abundance across European countries from 1990 to 2023. Unlike previous large-scale studies that treated migratory strategy as a species-level trait (e.g., Howard et al. 2020; Burns et al. 2021; Vickery et al. 2023), we took into account variation in migratory propensity both among and within populations. By integrating multiple complementary data sources, we provide a spatially explicit characterisation of migratory strategies for European landbirds at the population (and sometimes sub-population) level, covering over 90% of national common landbird abundance in most of the countries analysed (Figure S3). Our analyses show that, in a context of overall declining abundance of common birds since 1990 (Figure S6), long-distance migrants have declined the most, followed by short-distance migrants, whereas residents have remained stable or increased except in Northern Europe. As a result, the proportion of migrants declined across Europe, mainly in Southern Europe, while remaining stable or increasing in the north.

### Patterns of migratory strategies across Europe

We found that the dominance of migratory birds generally increases along a South-West to North-East gradient across Europe (Figure 1). This pattern is consistent with macroecological studies showing that migratory behaviour is more prevalent in highly seasonal environments, as found in high latitudes and continental climates (Somveille et al. 2013, 2015). Our results refine this picture by distinguishing between migratory strategies. We found the same broad gradient for short-distance migrants (Figure 1b), however for long-distance migrants we obtained an hourglass-shaped pattern (Figure 1a): they are highly abundant in Northern communities (almost 40% of total abundance in Fennoscandia), at their lowest in Central and Western Europe (e.g., 6% in Austria) and in the British Isles (7% in the UK), but relatively abundant in Southern countries (e.g., 17% in Spain, 14% in Italy). Overall, our results indicate that Fennoscandia is strongly dominated by migrants (both short and long-distance), Central Europe by residents and short-distance migrants (as also found by Storch et al. 2023), and the British Isles and Southern Europe by residents (up to 83% in Spain and in the UK).

These patterns should be interpreted with caution. We used ringing recoveries to distinguish resident from short-distance migrants within national populations, and ringing effort varied substantially across the study area. Although we corrected the proportions of recoveries for ringing effort, and our main results were robust to estimates derived without correction (Figure S14–S16), the greatest uncertainty remains for regions with little ringing data (Figure S2). Indeed, if a national population spending the winter in a given region was not detected by ringing recoveries, then that region was not identified as part of its wintering range, leading to the overestimation of the relative importance of the other wintering regions where the population was detected. Across species, this likely resulted in an underestimation of the importance of short-distance migration, particularly for populations migrating eastwards or towards North Africa, and a corresponding overestimation of the importance of residency. This may contribute to explaining the low prevalence of short-distance migrants in southern countries (e.g., Portugal, Slovenia, Greece), which may result from the non-detection of short-distance migration into Northern Africa. In addition, ringed birds are more likely to be recovered near their ringing site than on distant wintering grounds, which further biases estimates towards residency, particularly in regions with high ringing effort such as the British Isles (Figure 1).

### Dynamics of European bird populations across migratory strategies

Our results confirm the overall ongoing decline of long-distance migrants across Europe between 1990 and 2023 (Figure 2), consistent with previous continental-scale studies (Burns et al. 2021; Vickery et al. 2023; Sanderson et al. 2006; Howard et al. 2020; Table 1). Such declines have been attributed to a higher exposure to cumulative threats across the annual cycle (Buchan et al., 2023; La Sorte et al., 2018; Tøttrup et al., 2012). In addition, species may potentially experience mismatches with the phenology of food resources under climate change (Robertson et al., 2024), although the evidence that this has population level consequences is still weak (e.g. Franks et al. 2018). Focusing on landbirds, we found that, over this period, long-distance migrants declined by nearly 30% in total abundance, and by 27% in multi-species index across Europe (Figure 2). The similarity in the results between these two metrics indicates that such declines affected a broad set of populations, rather than being driven by just a few abundant ones. This said, population trends did vary among migratory populations, including many that declined (e.g. Barn Swallow *Hirundo rustica* in Bulgaria) and some that increased (e.g., Barn Swallow in the Netherlands) (Figure S9).

As we predicted, short-distance migrants declined less than long-distance migrants, but the magnitude of their decline differed markedly when measured through the multi-species index (4.5% loss across Europe) or total abundance (10%) (Figure 2). This contrast indicates that the declines in total abundance were driven by a few numerically dominant populations, such as the European Robin *Erithacus rubecula* in Finland or the Chaffinch *Fringilla coelebs* in Estonia (Figure S9), rather than by consistent declines across all species. This pattern mirrors broader trends in European bird communities, where common species have declined more strongly than rarer ones (Inger et al. 2015; Burns et al. 2021).

In contrast, residents (which account for about half of the total European population size), remained broadly stable, with a small increase in their total abundance (3%). This is in line with the results obtained by Vickery et al. (2023) and Sanderson et al. (2006) but not with Burns et al. (2021), who found declines of over 20% in resident populations. This contrast is likely to result (at least partially) from a difference in the period covered by the analysis: 1980–2017 versus 1990–2023. Indeed, Rigal et al. (2023) found that breeding birds across Europe (all migratory strategies combined) declined substantially in total abundance between 1980 and the mid-1990s, before stabilising.

Given that in our study we have characterised the trends for each migratory strategy in finer detail than in previous analyses, accounting for variation in migratory behaviour across and within populations, rather than assigning a single strategy to each species, we are confident that our results provide a more accurate picture of the observed trajectories than previous studies, for the period and species we covered.

This said, our results should be interpreted in light of a key assumption we needed to make (given data availability) for populations with mixed migratory strategies: that migratory and resident sub-populations follow the same demographic trajectory as the national population, and hence the proportion of individuals following each strategy remained constant throughout the study period. If instead migratory sub-populations have declined more rapidly than resident ones (likely, given our results), or if conversely the proportion of residents has increased because populations have shifted towards greater winter residency (Van Vliet et al., 2009) and shorter migration distances over time (Visser et al., 2009), then our results are conservative, with the relative declines of migratory birds relative to residency being even stronger in reality than in our estimates.

Despite the overall decline in migratory birds, our models suggest that the rate of decline in total abundance of both long- and short-distance migrants has slowed in recent years (Figure 2b). This deceleration is not apparent in the multi-species indices (Figure 2a), which may indicate that the most abundant populations are now declining less rapidly than rarer ones, and/or that some migratory populations are now increasing (e.g., Eurasian Blackcap in Denmark). Whether this pattern reflects an early sign of recovery remains uncertain. For instance, rainfall in the Sahel has partially recovered since the mid-1990s, which has been linked to population increases in some trans-Saharan migrants such as the Common Whitethroat *Curruca communis* and Common Redstart *Phoenicurus phoenicurus* (Zwarts et al., 2023), although the extent to which this recovery benefits migratory birds more broadly remains unclear. Conservation measures may also contribute to recovery, including hunting moratoria as shown for the European Turtle Dove *Streptopelia turtur*, within the European Union (Carboneras et al., 2024).

The combined effect of all these changes across different migratory strategies was an overall decline in European bird communities over the study period (1990-2023) of 14% when measured through the multi-species index and of 9% in total abundance (Figure S6). The magnitude of this decline is smaller than previous estimates reporting reductions of around 17-25% in European bird abundance between 1980 and the mid-2010s (Burns et al. 2021; Rigal et al. 2023), which may indicate that the rate of decline of European bird abundance has slowed in recent years, although it may also reflect differences in the set of species analysed. In any case, our results show that these changes are happening alongside a restructuring of avian communities as they become increasingly dominated by resident species both at the country (Figure 5) and at the continental levels (from 47.2% to 53.2% residents).

### Spatial variation in migratory community trajectories

While confirming that bird communities across Europe are undergoing significant changes to their composition in terms of migratory strategies, our study shows that these continental-scale results mask contrasting spatial patterns between countries, with geographical variation across migratory strategies.

The steepest declines in both total abundance and multi-species indices were observed in Northern Europe, affecting all three migratory strategies (Figures 3-4). For long-distance migrants, these results are consistent with previous studies in Estonia (Kuresoo et al. 2011) and Finland (Laaksonen and Lehikoinen 2013). In Northern countries, these declines are partly driven by the trajectories of highly abundant species. In particular, the Willow Warbler accounted for over half of all long-distance migrants in Sweden and Finland in 1990, where it declined by 38% and 34%, respectively, during the study period (Figure S9), thus single-handedly affecting overall trends in the total abundance of long-distance migrants in those countries (Figures 4, S8). Northern communities also contain a higher proportion of migratory species, increasing their exposure to climatic and environmental disruptions affecting phenology, habitat quality, and resource availability along migratory routes (Zurell et al., 2018). However, declines in the north were not restricted to migrants or to the well-documented decline of farmland birds (Laaksonen and Lehikoinen 2013; Rigal et al. 2023): our results indicate that entire bird communities are declining, including resident forest species such as Willow Tit *Poecile montanus* (70% decline in Sweden and Finland) and Goldcrest *Regulus regulus* (>60% decline in Estonia; Figure S9). Such declines may be masked in aggregated forest bird indicators, which have remained broadly stable in Norway and Sweden (Rigal and Knape 2024). These patterns likely reflect multiple interacting drivers, such as intensive forestry practices and potentially climate warming at high latitudes leading to greater vulnerability of cold-dwelling species (Howard et al. 2015; Virkkala et al. 2008; Cours et al. 2025; Lehikoinen et al. 2021; Tayleur et al. 2016).

In contrast, in Central and Southern Europe the trends for resident and short-distance migratory species (but not for long-distance migrants) were generally more stable, or even positive. This echoes results from national studies such as Bowler et al. (2021), who found that migratory species experienced a steeper decline in abundance than resident species in the Czech Republic and Switzerland. Likewise, Kamp et al. (2021), reported declines in abundance among long-distance migrants but relative stability of short-distance migrants in Germany, while Tirozzi et al. (2021) found that in Italy both short- and long-distance migrants declined, while residents increased. Although differences in time periods and the set of species considered complicate direct comparisons between these national studies, our analysis provides a harmonised continental perspective on changes in migratory composition of national communities.

The stable or increasing trends observed among resident species in Central and Southern Europe may partly reflect the influence of climate warming, as milder winters can improve the survival of non-migratory species (Pearce-Higgins et al., 2015). Nonetheless, short-distance migrants often showed contrasting trajectories between neighbouring countries, for example, increasing in Hungary while declining in Austria (Figure 4). Previous studies have also reported significant geographical variation in population trends across Europe, both between species groups (Bowler et al. 2021) and within species, such as the Willow Warbler in the British Isles (Morrison et al., 2010). This variability likely reflects the importance of regional or national drivers, including differences in climate and land-use change, such as agricultural intensification and forestry practices, which affect species according to their ecological requirements (Rigal et al., 2023).

Finally, compiling our dataset revealed gaps in knowledge of bird movements. We had to exclude three South-Eastern countries (Greece, Romania and Bulgaria) from the national-scale analyses because data coverage on population trends was insufficient, particularly for short-distance migrants. Furthermore, ringing effort is unevenly distributed across Europe and North Africa (Figure S2), so movements of birds wintering in North Africa or Eastern Europe remain poorly documented. Broader coverage of ringing and tracking efforts in these regions will be needed to describe European migratory movements comprehensively, link them to population dynamics and assess changes in European bird communities. In addition, national population sizes, largely reported under the EU Birds Directive, are estimated using different methods across countries, which may limit comparisons of absolute abundance between countries, although this does not affect temporal trends within countries. In particular, population sizes reported for Spain in the ERLoB stand out, representing 30% of the total estimated abundance of landbirds of the countries included in the study (Figure S4 and S5). Including Spain did not change our overall results and mainly strengthened the increase in total abundance of residents relative to the other strategies (Figures S4 and S10), but it calls for caution when interpreting absolute abundance estimates.

These limitations notwithstanding, our results indicate that the reorganisation of European migratory bird communities is uneven across the continent (Figure 5), with a broad gradient towards an increasing dominance of resident species at lower latitudes. However, this pattern conceals however important differences in the underlying processes: in Central and Southern Europe, it primarily reflects stronger declines among migratory species, whereas in Northern Europe the relatively stable proportion of migrants is the result of comparable declines affecting both migratory and resident species.

## Conclusion

Our results show that European breeding bird communities are being progressively reshaped according to migratory strategy. Long-distance migrants are undergoing substantial declines, short-distance migrants show intermediate trends, and resident species remain broadly stable. These divergent trajectories are not evenly distributed across the continent, with the strongest declines occurring in Northern Europe.

This growing imbalance between migratory strategies signals a general shift toward increasingly resident-dominated bird communities. While our results provide some insights into the potential drivers of these changes, identifying their precise causes is beyond the scope of our study. The mechanisms underlying these trends are likely complex and interacting, reflecting both differential exposure to environmental pressures (such as climate change and land-use change) and differences in life-history traits that shape species’ vulnerability. Important next steps will be to understand how this restructuring interacts with other well-documented changes in European bird communities, such as the decline of farmland birds, and to link trends to more detailed information on wintering areas (e.g., West, Central and East Africa or Asia for long-distance migrants; Western, Southern and Eastern Europe for short-distance migrants) and on habitat preferences (e.g., woodland, open and semi-open habitats, or wetlands).

The dataset presented here provides a unique and up-to-date framework for investigating these processes. By integrating population trends, abundance estimates, and migratory behaviour across countries, our study offers the first continental-scale quantification of the ongoing restructuration of European bird communities according to migratory strategies. These compositional changes may have important ecological consequences, given the key role migratory birds play in linking distant ecosystems and maintaining ecosystem functions (Bauer and Hoye 2014). Maintaining and expanding coordinated efforts to monitor population size (through PECBMS) and understand the movements of migratory birds (e.g., through ringing and the growing European network of Motus receivers; Taylor et al. 2017), will be essential to better understand, and ultimately mitigate, the cascading effects of global changes on migratory birds and the ecosystems they connect.

## Acknowledgments

From the song of a bird recorded locally to continental-scale indicators, this study would not have been possible without an extensive network of data collection and knowledge sharing. We are deeply grateful to the thousands of volunteers across Europe who contribute to bird monitoring programmes and whose efforts are essential for advancing our understanding of bird population dynamics and movements.

We also thank the national coordinators of the Common Bird Monitoring Schemes for organising surveys, maintaining long-term monitoring efforts, and ensuring the quality of national datasets. In particular, we acknowledge the coordinators of the Pan-European Common Bird Monitoring Scheme (PECBMS) for their substantial work in compiling, harmonising, and maintaining bird monitoring data at the continental scale: Benjamin Seaman, Norbert Teufelbauer (**Austria**), Antoine Derouaux, Olivier Dochy, Thierry Onkelinx, Marius Pailhès, Jean-Yves Paquet (**Belgium**), Iordan Hristov, Georgi Popgeorgiev (**Bulgaria**), Martin Hellicar, Christina Ieronymidou, Stalo Leontiou (**Cyprus**), Alena Jechumtál Skálová, Jiří Reif, Zdeněk Vermouzek, Petr Voříšek (**Czechia**), Daniel Palm Eskildsen (**Denmark**), Meelis Leivits, Renno Nellis (**Estonia**), Heikki Helle, Aleksi Lehikoinen (**Finland**), Benoît Fontaine, Victoire Kuhn (**France**), Malte Busch, Martin Flade, Bettina Gerlach, Johannes Schwarz, Sven Trautmann (**Germany**), Aris Manolopoulos, Danae Portolou (**Greece**), Balázs Csibrány, Károly Nagy, Tibor Szép (**Hungary**), Mattia Brambilla, Gianpiero Calvi, Simonetta Cutini, Matteo Fontanella, Federica Luoni, Roberta Righini, Laura Silva (**Italy**), Ainārs Auniņš, Oskars Keišs, Ieva Mārdega (**Latvia**), Petras Kurlavičius, Renata Mackevičienė, Gintaras Riauba (**Lithuania**), Gilles Biver, Katharina Klein, Cindy Redel (**Luxembourg**), Arjan Boele, Chiel Boom, Joost van Bruggen, Adriaan Gmelig Meyling, Kees Koffijberg, Dorine Jansen, Jan-Willem Vergeer, Richard Verweij, Dirk Zoetebier (**the Netherlands**), Knut Eie, John Atle Kålås, Torstein Myhre, Ingar Jostein Øien, Tom Roger Østerås, Diego Pavón Jordán, Paul Shimmings, Bård Gunnar Stokke, Roald Vang (**Norway**), Tomasz Chodkiewicz, Przemysław Chylarecki, Paulina Hapon, Łukasz Wardecki (**Poland**), Hany Alonso, Rúben Coelho, Cátia Gouveia, Guillaume Réthoré (**Portugal**), Brian Burke, Sinéad Cummins, John Kennedy, Lesley Lewis, Andrew Lynch, Alyn Walsh (**the Republic of Ireland**), Cristi Domşa, Judit Veres-Szászka, Anca Pui (**Romania**), Peter Lešo, Jozef Ridzoň, Soňa Svetlíková (**Slovakia**), Dejan Bordjan, Matej Gamser, Primož Kmecl, Davide Scridel (**Slovenia**), Marc Anton, Lluís Brotons, Virginia Escandell, Emilio Escudero, David Garcia, Sergi Herrando, David Martí, Juan Carlos del Moral (**Spain**), Martin Green, Fredrik Haas, Åke Lindström, Martin Stjernman (**Sweden**), Roman Bühler, Christian Rogenmoser, Nicolas Strebel, Samuel Wechsler (**Switzerland**), Caroline Brighton, James Heywood, Dario Massimino, David Noble (**the United Kingdom**).

We also thank those who submitted data on bird population sizes and trends across the EU and the national coordinators.

## Declaration of Generative AI and AI-assisted technologies in the writing process

During the preparation of this work, the authors used Deepl to support the writing. They have reviewed and edited the content and assume full responsibility for the content of the published article.

## Data, scripts, code, and supplementary information availability

Table S2, listing the populations included in the analyses, with their migratory strategy (or, for partially migratory populations, the proportion of individuals assigned to each strategy) and their estimated change in population size over the period of monitoring, is available at https://doi.org/10.5281/zenodo.24284226

All other data, code and supplementary materials are available at https://doi.org/10.5281/zenodo.24322562

## Bibliography

Bauer, S., & Hoye, B. J. (2014). Migratory animals couple biodiversity and ecosystem functioning worldwide. Science (New York, N.Y.), 344(6179), 1242552. 10.1126/science.1242552

Billerman, S. M., Keeney, B. K., Kirwan, G. M., Medano, F., Sly, N. D., & Smith, M. G. (2025). Birds of the world. Cornell Lab of Ornithology. https://birdsoftheworld.org/bow/home

BirdLife International. (2021). European Red List of Birds.

BirdLife International. (2026). State of the world’s birds: A focus on migratory species and flyway. https://datazone.birdlife.org/publications/state-of-the-worlds-birds-2026

BirdLife International and Handbook of the Birds of the World. (2024). Bird species’ distribution maps of the world. https://datazone.birdlife.org/contact-us/request-our-data.

Bogaart, P., Loo M. van der, Pannekoek, J., & Netherlands, S. (2026). rtrim: Trends and Indices for Monitoring Data (Version 2.3.1) [Computer software]. https://cran.r-project.org/web/packages/rtrim/index.html

Bowler, D., Richter, R. L., Eskildsen, D., Kamp, J., Moshøj, C. M., Reif, J., Strebel, N., Trautmann, S., & Voříšek, P. (2021). Geographic variation in the population trends of common breeding birds across central Europe. Basic and Applied Ecology, 56, 72–84. 10.1016/j.baae.2021.07.004

Briedis, M., Bauer, S., Adamík, P., Alves, J. A., Costa, J. S., Emmenegger, T., Gustafsson, L., Koleček, J., Krist, M., Liechti, F., Lisovski, S., Meier, C. M., Procházka, P., & Hahn, S. (2020). Broad-scale patterns of the Afro-Palaearctic landbird migration. Global Ecology and Biogeography, 29(4), 722–735. 10.1111/geb.13063

Buchan, C., Gilroy, J. J., Catry, I., Hewson, C. M., Atkinson, P. W., & Franco, A. M. A. (2023). Combining remote sensing and tracking data to quantify species’ cumulative exposure to anthropogenic change. Global Change Biology, 29(23), 6679–6692. 10.1111/gcb.16974

Burns, F., Eaton, M. A., Burfield, I. J., Klvaňová, A., Šilarová, E., Staneva, A., & Gregory, R. D. (2021). Abundance decline in the avifauna of the European Union reveals cross-continental similarities in biodiversity change. Ecology and Evolution, 11(23), 16647–16660. 10.1002/ece3.8282

Carboneras, C., Šilarová, E., Škorpilová, J., & Arroyo, B. (2024). Rapid population response to a hunting ban in a previously overharvested, threatened landbird. Conservation Letters, 17(6), e13057. 10.1111/conl.13057

Chapman, B. B., Brönmark, C., Nilsson, J.-Å., & Hansson, L.-A. (2011). The ecology and evolution of partial migration. Oikos, 120(12), 1764–1775. 10.1111/j.1600-0706.2011.20131.x

Collar, N., & Christie, D. (2020). Eurasian Blackbird (Turdus merula), version 1.0. Birds of the World. 10.2173/bow.eurbla.01

Copernicus Climate Change Service. (2023). ERA5 monthly averaged data on single levels from 1940 to present. Climate Data Store (CDS). 10.24381/cds.f17050d7

Cours, J., Elo, M., Pithon, J., Triviño, M., Mönkkönen, M., Hagge, J., Lehikoinen, A., & Duflot, R. (2025). Changes in abundance and distribution of European forest bird populations depend on biome, ecological specialisation and traits. Ecography, 2025(7), e07582. 10.1111/ecog.07582

Cuchot, P., Bonnet, T., Robinson, R. A., Arizaga, J., Fiedler, W., Geiter, O., Henshaw, I., Herrmann, C., van der Jeugd, H., Karcza, Z., Leal, A., Lehikoinen, P., von Rönn, J. A. C., Thorup, K., Teplitsky, C., & Henry, P.-Y. (2025). Phenological Plasticity and Its Temperature-Related Drivers in Common Songbirds Across Europe. Global Change Biology, 31(11), e70600. 10.1111/gcb.70600

Delmore, K. E., Van Doren, B. M., Conway, G. J., Curk, T., Garrido-Garduño, T., Germain, R. R., Hasselmann, T., Hiemer, D., van der Jeugd, H. P., Justen, H., Lugo Ramos, J. S., Maggini, I., Meyer, B. S., Phillips, R. J., Remisiewicz, M., Roberts, G. C. M., Sheldon, B. C., Vogl, W., & Liedvogel, M. (2020). Individual variability and versatility in an eco-evolutionary model of avian migration. Proceedings of the Royal Society B: Biological Sciences, 287(1938), 20201339. 10.1098/rspb.2020.1339

Devictor, V., Julliard, R., Couvet, D., & Jiguet, F. (2008). Birds are tracking climate warming, but not fast enough. Proceedings of the Royal Society B: Biological Sciences, 275(1652), 2743–2748. 10.1098/rspb.2008.0878

Doxa, A., Bas, Y., Paracchini, M. L., Pointereau, P., Terres, J., & Jiguet, F. (2010). Low-intensity agriculture increases farmland bird abundances in France. Journal of Applied Ecology, 47(6), 1348–1356. 10.1111/j.1365-2664.2010.01869.x

Du Feu, C. R., Clark, J. A., Schaub, M., Fiedler, W., & Baillie, S. R. (2016). The EURING Data Bank – a critical tool for continental-scale studies of marked birds. Ringing & Migration, 31(1), 1–18. 10.1080/03078698.2016.1195205

Fattorini, N., Costanzo, A., Romano, A., Rubolini, D., Baillie, S., Bairlein, F., Spina, F., & Ambrosini, R. (2023). Eco-evolutionary drivers of avian migratory connectivity. Ecology Letters, 26(7), 1095–1107. 10.1111/ele.14223

Fontaine, B., Moussy, C., Chiffard, J., Dupuis, J., Corolleur, E., Schmaltz, L., Lorrilliere, R., Loïs, G., & Gaudard, C. (2021). Suivi des oiseaux communs en France 1989-2019: 30 ans de suivis participatifs.

Gaston, K. J. (2010). Valuing Common Species. Science, 327(5962), 154–155. 10.1126/science.1182818

Howard, C., Stephens, P. A., Pearce-Higgins, J. W., Gregory, R. D., Butchart, S. H. M., & Willis, S. G. (2020). Disentangling the relative roles of climate and land cover change in driving the long-term population trends of European migratory birds. Diversity and Distributions, 26(11), 1442–1455. 10.1111/ddi.13144

Howard, C., Stephens, P. A., Pearce-Higgins, J. W., Gregory, R. D., & Willis, S. G. (2015). The drivers of avian abundance: Patterns in the relative importance of climate and land use. Global Ecology and Biogeography, 24(11), 1249–1260. 10.1111/geb.12377

Inger, R., Gregory, R., Duffy, J. P., Stott, I., Voříšek, P., & Gaston, K. J. (2015). Common European birds are declining rapidly while less abundant species’ numbers are rising. Ecology Letters, 18(1), 28–36. 10.1111/ele.12387

Kamp, J., Frank, C., Trautmann, S., Busch, M., Dröschmeister, R., Flade, M., Gerlach, B., Karthäuser, J., Kunz, F., Mitschke, A., Schwarz, J., & Sudfeldt, C. (2021). Population trends of common breeding birds in Germany 1990–2018. Journal of Ornithology, 162(1), 1–15. 10.1007/s10336-020-01830-4

Kirby, J. S., Stattersfield, A. J., Butchart, S. H. M., Evans, M. I., Grimmett, R. F. A., Jones, V. R., O’Sullivan, J., Tucker, G. M., & Newton, I. (2008). Key conservation issues for migratory land- and waterbird species on the world’s major flyways. Bird Conservation International, 18(S1), S49–S73. 10.1017/S0959270908000439

Kroodsma, D. E., Brewer, D., Christie, D., & Marks, J. S. (2020). Eurasian Wren (Troglodytes troglodytes), version 1.0. Birds of the World. 10.2173/bow.winwre4.01

Kuemmerle, T., Levers, C., Erb, K., Estel, S., Jepsen, M. R., Müller, D., Plutzar, C., Stürck, J., Verkerk, P. J., Verburg, P. H., & Reenberg, A. (2016). Hotspots of land use change in Europe. Environmental Research Letters, 11(6), 064020. 10.1088/1748-9326/11/6/064020

Kuresoo, A., Pehlak, H., & Nellis, R. (2011). Population trends of common birds in Estonia in 1983–2010; pp. 88–110.Estonian Journal of Ecology, 60(2), 88–110. 10.3176/eco.2011.2.02

La Sorte, F. A., Fink, D., & Johnston, A. (2018). Seasonal associations with novel climates for North American migratory bird populations. Ecology Letters, 21(6), 845–856. 10.1111/ele.12951

Laaksonen, T., & Lehikoinen, A. (2013). Population trends in boreal birds: Continuing declines in agricultural, northern, and long-distance migrant species. Biological Conservation, 168, 99–107. 10.1016/j.biocon.2013.09.007

Lampila, P., Mönkkönen, M., & Desrochers, A. (2005). Demographic Responses by Birds to Forest Fragmentation. Conservation Biology, 19(5), 1537–1546. 10.1111/j.1523-1739.2005.00201.x

Lehikoinen, A., Lindström, Å., Santangeli, A., Sirkiä, P. M., Brotons, L., Devictor, V., Elts, J., Foppen, R. P. B., Heldbjerg, H., Herrando, S., Herremans, M., Hudson, M.-A. R., Jiguet, F., Johnston, A., Lorrilliere, R., Marjakangas, E.-L., Michel, N. L., Moshøj, C. M., Nellis, R., … van Turnhout, C. (2021). Wintering bird communities are tracking climate change faster than breeding communities. Journal of Animal Ecology, 90(5), 1085–1095. 10.1111/1365-2656.13433

Marra, P. P., Cooper, N. W., Loss, S. R., Hunter, D., Bravo Hernández, E., Citegetse, G., Hewson, C. M., Jahn, A. E., Jones, V. R., Ndang’ang’a, P. K., Ozsanlav-Harris, L., Rosenberg, K. V., Vickery, J. A., Viswanathan, A., & Yong, D. L. (2026). Status, threats and conservation of Earth’s migratory birds. Nature Reviews Biodiversity, 2(8), 488–505. 10.1038/s44358-026-00177-7

Møller, A. P., Rubolini, D., & Lehikoinen, E. (2008). Populations of migratory bird species that did not show a phenological response to climate change are declining. Proceedings of the National Academy of Sciences, 105(42), 16195–16200. 10.1073/pnas.0803825105

Monnet, A.-C., Cairo, M., Deguines, N., Jiguet, F., Vimont, M., Fontaine, C., & Porcher, E. (2026). Common birds have higher abundances in croplands with lower pesticide purchases. Proceedings of the Royal Society B: Biological Sciences, 293(2062), 20252370. 10.1098/rspb.2025.2370

Morrison, C. A., Robinson, R. A., Clark, J. A., & Gill, J. A. (2010). Spatial and temporal variation in population trends in a long-distance migratory bird. Diversity and Distributions, 16(4), 620–627. 10.1111/j.1472-4642.2010.00663.x

Newton, I. (2023). The Migration Ecology of Birds (2nd edn). Elsevier/Acad. Press. https://shop.elsevier.com/books/the-migration-ecology-of-birds/newton/978-0-12-823751-9

Pannekoek, J., & van Strien, A. (2005). TRIM 3 Manual (TRends & Indices for Monitoring data).

Pearce-Higgins, J. W., Eglington, S. M., Martay, B., & Chamberlain, D. E. (2015). Drivers of climate change impacts on bird communities. Journal of Animal Ecology, 84(4), 943–954. 10.1111/1365-2656.12364

R Core Team. (2023). R: A Language and Environment for Statistical Computing (Version 4.3.2) [Computer software]. R Foundation for Statistical Computing. https://www.R-project.org/

Rigal, S., Dakos, V., Alonso, H., Auniņš, A., Benkő, Z., Brotons, L., Chodkiewicz, T., Chylarecki, P., de Carli, E., del Moral, J. C., Domşa, C., Escandell, V., Fontaine, B., Foppen, R., Gregory, R., Harris, S., Herrando, S., Husby, M., Ieronymidou, C., … Devictor, V. (2023). Farmland practices are driving bird population decline across Europe. Proceedings of the National Academy of Sciences, 120(21), e2216573120. 10.1073/pnas.2216573120

Rigal, S., Devictor, V., & Dakos, V. (2020). A method for classifying and comparing non-linear trajectories of ecological variables. Ecological Indicators, 112, 106113. 10.1016/j.ecolind.2020.106113

Rigal, S., & Knape, J. (2024). Investigating the heterogeneity within Wild bird indices in Europe. Biological Conservation, 290, 110452. 10.1016/j.biocon.2024.110452

Robertson, E. P., La Sorte, F. A., Mays, J. D., Taillie, P. J., Robinson, O. J., Ansley, R. J., O’Connell, T. J., Davis, C. A., & Loss, S. R. (2024). Decoupling of bird migration from the changing phenology of spring green-up. Proceedings of the National Academy of Sciences, 121(12), e2308433121. 10.1073/pnas.2308433121

Rosenberg, K. V., Dokter, A. M., Blancher, P. J., Sauer, J. R., Smith, A. C., Smith, P. A., Stanton, J. C., Panjabi, A., Helft, L., Parr, M., & Marra, P. P. (2019). Decline of the North American avifauna. Science, 366(6461), 120–124. 10.1126/science.aaw1313

Runge, C. A., Martin, T. G., Possingham, H. P., Willis, S. G., & Fuller, R. A. (2014). Conserving mobile species. Frontiers in Ecology and the Environment, 12(7), 395–402. 10.1890/130237

Sanderson, F. J., Donald, P. F., Pain, D. J., Burfield, I. J., & van Bommel, F. P. J. (2006). Long-term population declines in Afro-Palearctic migrant birds. Biological Conservation, 131(1), 93–105. 10.1016/j.biocon.2006.02.008

Soldaat, L. L., Pannekoek, J., Verweij, R. J. T., Van Turnhout, C. A. M., & Van Strien, A. J. (2017). A Monte Carlo method to account for sampling error in multi-species indicators. Ecological Indicators, 81, 340–347. 10.1016/j.ecolind.2017.05.033

Somveille, M., Manica, A., Butchart, S. H. M., & Rodrigues, A. S. L. (2013). Mapping Global Diversity Patterns for Migratory Birds. PLOS ONE, 8(8), e70907. 10.1371/journal.pone.0070907

Somveille, M., Rodrigues, A. S. L., & Manica, A. (2015). Why do birds migrate? A macroecological perspective. Global Ecology and Biogeography, 24(6), 664–674. 10.1111/geb.12298

Spina, F., Baillie, S. R., Bairlein, F., Fiedler, W., & Thorup, K. (2022). Eurasian African Bird Migration Atlas. EURING/CMS. https://migrationatlas.org/

Stephens, P. A., Mason, L. R., Green, R. E., Gregory, R. D., Sauer, J. R., Alison, J., Aunins, A., Brotons, L., Butchart, S. H. M., Campedelli, T., Chodkiewicz, T., Chylarecki, P., Crowe, O., Elts, J., Escandell, V., Foppen, R. P. B., Heldbjerg, H., Herrando, S., Husby, M., … Willis, S. G. (2016). Consistent response of bird populations to climate change on two continents. Science, 352(6281), 84–87. 10.1126/science.aac4858

Storch, D., Koleček, J., Keil, P., Vermouzek, Z., Voříšek, P., & Reif, J. (2023). Decomposing trends in bird populations: Climate, life histories and habitat affect different aspects of population change. Diversity and Distributions, 29(4), 572–585. 10.1111/ddi.13682

Tayleur, C. M., Devictor, V., Gaüzère, P., Jonzén, N., Smith, H. G., & Lindström, Å. (2016). Regional variation in climate change winners and losers highlights the rapid loss of cold-dwelling species. Diversity and Distributions, 22(4), 468–480. 10.1111/ddi.12412

Taylor, P., Crewe, T., Mackenzie, S., Lepage, D., Aubry, Y., Crysler, Z., Finney, G., Francis, C., Guglielmo, C., Hamilton, D., Holberton, R., Loring, P., Mitchell, G., Norris, D. R., Paquet, J., Ronconi, R., Smetzer, J., Smith, P., Welch, L., & Woodworth, B. (2017). The Motus Wildlife Tracking System: A collaborative research network to enhance the understanding of wildlife movement. Avian Conservation and Ecology, 12(1). 10.5751/ACE-00953-120108

Tirozzi, P., Orioli, V., Dondina, O., Kataoka, L., & Bani, L. (2021). Species Traits Drive Long-Term Population Trends of Common Breeding Birds in Northern Italy. Animals, 11(12), 3426. 10.3390/ani11123426

Tøttrup, A. P., Klaassen, R. H. G., Kristensen, M. W., Strandberg, R., Vardanis, Y., Lindström, Å., Rahbek, C., Alerstam, T., & Thorup, K. (2012). Drought in Africa Caused Delayed Arrival of European Songbirds. Science, 338(6112), 1307–1307. 10.1126/science.1227548

Van Vliet, J., Musters, C. J. M., & Ter Keurs, W. J. (2009). Changes in migration behaviour of Blackbirds Turdus merula from the Netherlands. Bird Study, 56(2), 276–281. 10.1080/00063650902792148

Vickery, J. A., Mallord, J. W., Adams, W. M., Beresford, A. E., Both, C., Cresswell, W., Diop, N., Ewing, S. R., Gregory, R. D., Morrison, C. A., Sanderson, F. J., Thorup, K., Van Wijk, R. E., & Hewson, C. M. (2023). The conservation of Afro-Palaearctic migrants: What we are learning and what we need to know? Ibis, 165(3), 717–738. 10.1111/ibi.13171

Virkkala, R., Heikkinen, R. K., Leikola, N., & Luoto, M. (2008). Projected large-scale range reductions of northern-boreal land bird species due to climate change. Biological Conservation, 141(5), 1343–1353. 10.1016/j.biocon.2008.03.007

Visser, M. E., Perdeck, A. C., Van BALEN, J. H., & Both, C. (2009). Climate change leads to decreasing bird migration distances. Global Change Biology, 15(8), 1859–1865. 10.1111/j.1365-2486.2009.01865.x

Yong, D. L., Heim, W., Chowdhury, S. U., Choi, C.-Y., Ktitorov, P., Kulikova, O., Kondratyev, A., Round, P. D., Allen, D., Trainor, C. R., Gibson, L., & Szabo, J. K. (2021). The State of Migratory Landbirds in the East Asian Flyway: Distributions, Threats, and Conservation Needs. Frontiers in Ecology and Evolution, 9. 10.3389/fevo.2021.613172

Zurell, D., Graham, C. H., Gallien, L., Thuiller, W., & Zimmermann, N. E. (2018). Long-distance migratory birds threatened by multiple independent risks from global change. Nature Climate Change, 8(11), Article 11. 10.1038/s41558-018-0312-9

Zwarts, L., Bijlsma, R. G., & Kamp J. van der. (2023). The Fortunes of Migratory Birds from Eurasia: Being on a Tightrope in the Sahel. Ardea, 111(1), 397–437. 10.5253/arde.2022.a29

